# Ultra-fast and highly sensitive protein structure alignment with segment-level representations and block-sparse optimization

**DOI:** 10.1101/2025.03.14.643159

**Authors:** Thomas Litfin, Yaoqi Zhou, Mark von Itzstein

## Abstract

Deep learning models for protein structure prediction have given rise to extreme growth in 3D structure data. As a result, traditional methods for geometric structure alignment are too slow to effectively search modern structure libraries. In this study we introduce SPfast – a fully geometric method for structure-based alignment which accelerates search by more than 2 orders of magnitude while increasing sensitivity by 21% and 5% compared with foldseek and TMalign respectively. Using the significant speed of SPfast to conduct more than 100B pairwise comparisons between *bona fide* uncharacterized proteins and a large-scale, annotated structure library uncovers new biological insights relating to type III secretion in pathogenic bacteria and identifies novel toxin-antitoxin systems. Putative SPfast-based functional assignments are supported by orthogonal evidence including shared genomic context and high-confidence AlphaFold3 complex modelling.

## Introduction

The protein function annotation pipeline relies on propagating experimentally validated annotations across closely related proteins. Historically, sequence profile alignment has been the gold-standard for identifying evolutionary relationships which suggest a shared functional role^1^. Sequence-based alignment is well suited to high-throughput search due to the extreme speed of the underlying algorithm – particularly when using heuristic approximations as implemented in methods such as BLAST^2^, HMMER3^3^ and MMSeqs2^4^. However, sequence similarity has limited sensitivity to detect functional relationships over great evolutionary distances leading to a large number of spurious ‘orphan’ genes without functional annotations^5^.

The AlphaFold protein structure database (AFDB)^6^ provides high quality model structures for more than 200M reference proteins catalogued in the UniProt^7^ database and the ESM metagenomic atlas^8^ contains an additional 772M structures predicted from metagenome sequencing projects. These large-scale protein structure libraries represent an opportunity to enhance the sensitivity of homology-based annotation using structure-based search. Prior works in this space have used highly sensitive geometric search to identify functional clues from the relatively small, PDB database^9^ or have sacrificed sensitivity by using tokenized structure alignments to search large model libraries^10–12^. Here we have developed new heuristics to enable practical high-throughput search using highly sensitive geometric alignments to enhance the quality of structure-based annotations.

Traditional approaches for geometric structure alignment utilize an iterative closest point (ICP) heuristic to identify paired correspondences between matched amino acids^13–16^. This heuristic involves an iterating process that alternates between superimposing structures, identifying an alignment and then updating the superposition to minimize root mean square deviation (RMSD) between aligned residues. However, alignments generated by the ICP algorithm are extremely sensitive to initial superpositions with early methods requiring manual assignment of matching residues to initialize a convergent alignment^13^. Modern methods provide an automated solution but rely on brute force enumeration of potential seeds derived primarily from contiguous peptide fragments^14–16^. In the worst case, this strategy generates a candidate set of initial alignments that scales quadratically with protein length (although many potential seeds are pruned based on fragment RMSD in practice). Similarly, the evaluation of each candidate alignment utilizes the intermolecular distance matrix as input to the Needleman-Wunch^17^ algorithm which also has nested quadratic complexity. While traditional tools such as TMalign^14^/USalign^16^ and SPalign^15^ have successfully leveraged this paradigm for pairwise alignment and PDB search, they cannot scale to handle the newfound abundance of predicted protein structure data.

A recently developed method, foldseek^18^, has been designed to alleviate the computational burden by representing proteins with an SE3-invariant structure-state alphabet and utilizing sequence-based acceleration heuristics in a one-pass alignment afforded by the MMSeqs2 framework^4^. Foldseek leverages a structure-state substitution matrix to characterize structural similarity and a *k*-mer based alignment prefilter coupled with highly optimized single instruction, multiple data (SIMD) intrinsics to accelerate database search. These heuristics facilitate lightning-fast execution times which enable practical search of large model databases. However, the acceleration is accompanied by a substantial reduction in search sensitivity – particularly when invoking the database prefilter – which may be prohibitive for applications that rely on exhaustive search.

In this work, we introduce SPfast – a significantly accelerated, fully geometric method for protein structure alignment that achieves state of the art search sensitivity with efficiency that can support high-throughput search. In SPfast we replace the traditional, Cα-based structure representation with idealized key points extracted from secondary structure segments. This coarse-grained, segment-level representation is used to generate a minimal set of alignment seeds by superimposing compatible pairs of idealized fragments. Candidate seeds are evaluated based on a segment-level alignment score which is used to screen potential structure matches. The top-ranking seed is finally used to guide a block-sparse, all-atom refinement of the pairwise alignment objective to produce a residue-level alignment. These heuristics are combined to increase the speed of geometric alignments by 2 orders of magnitude and up to 3 orders of magnitude when combined with a foldseek-based prefilter.

## Results

### Fold recognition of annotated SCOPe domains

Domain-level search sensitivity was evaluated by conducting an all-against-all comparison of 11,211 domains from the SCOPe dataset^19^ and ranking structures based on pairwise alignment scores using each method (Figure 1A). Of the existing methods in the literature, ranking structures based on an exhaustive optimization of SPscore (SPalign) achieved the highest mean sensitivity at first false positive across all 3 levels of the protein structure hierarchy (0.590 at the superfamily level). However, SPalign is impractical for use at scale – requiring 66.5 hours to complete the SCOPe benchmark. By comparison foldseek completed the benchmark in only 3.5 minutes on the same hardware – albeit at a considerable drop in search sensitivity (0.487). We found that the sensitivity of SPfast was equivalent to SPalign at the superfamily level (0.590), and superior to all other methods (including TMalign with sensitivity of 0.562), while completing the benchmark in just 37.3 minutes. SPfast re-ranking of structures identified by the foldseek prefilter (foldseekSP) was found to recover most of the performance of SPfast alone (0.557) and completed the benchmark in only 8.25 minutes (compared with just under 2 hours for foldseekTM to achieve sensitivity of 0.548).

**Figure 1.**
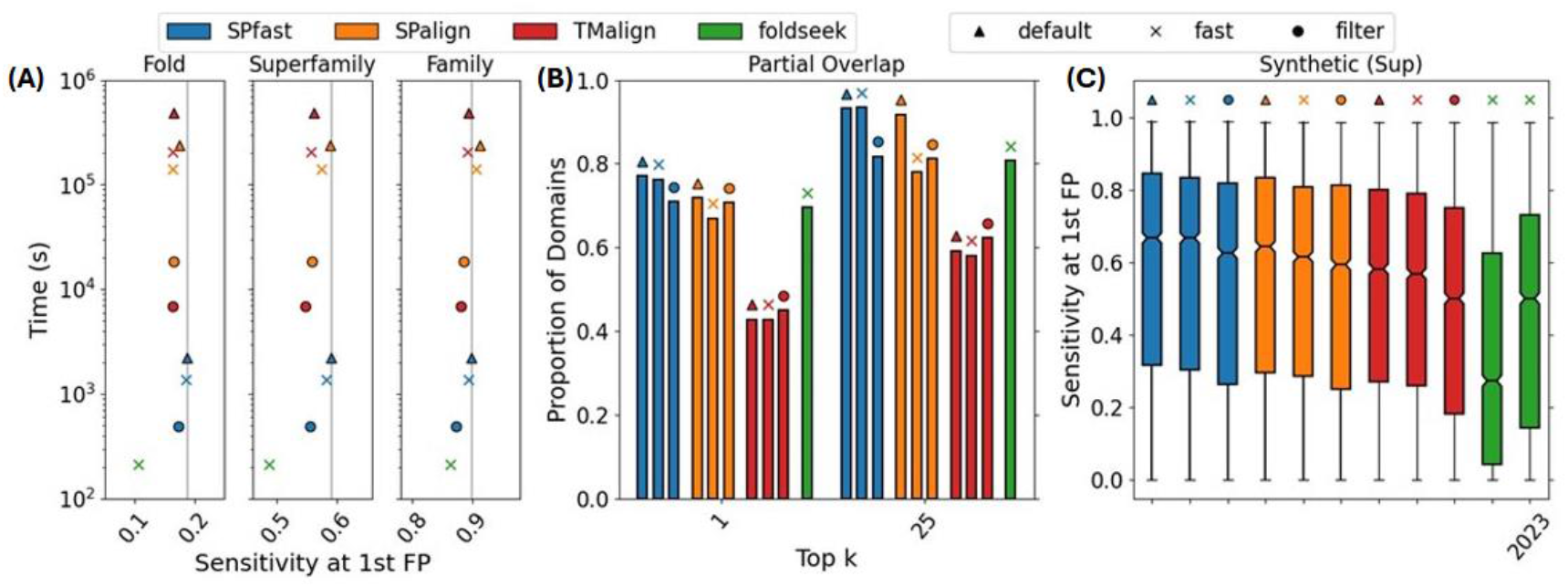
(A) Mean sensitivity at the first false positive (FP) for SCOPe domains at the fold, superfamily and family level and corresponding execution time for 11,211 x 11,211 all-by-all comparisons on an Intel Xeon E5-2670 @ 2.60GHz 16-core CPU. (B) Proportion of multi-domain AFDB proteins for which another protein with a single domain overlap can be identified in the top-k ranks. (C) Superfamily-level sensitivity at the first FP for synthetic SCOPe domains generated by RFdiffusion partial denoising (10 steps) and searched against natural SCOPe domains.

The optimization of SPfast alignment heuristics offered a continuous trade-off between search sensitivity and throughput. To investigate this trade-off, we evaluated the performance of SPfast by optimizing an SPscore objective and varying individual heuristic parameters (Supplementary Figure S1). The secondary structure prefilter had the largest impact on method performance. However, even the most stringent filtering criterion maintained superior sensitivity to foldseek (0.524 superfamily-level sensitivity at first false positive) and reduced SPfast execution time to just 17.5 minutes. Relaxing the ICP convergence criterion to 9% had an extremely mild impact on search sensitivity (0.567 c.f. 0.568 for the default 5%) and trimmed execution time to 35.8 minutes. Similarly, a segment-level alignment score cutoff of 5.5 reduced execution time to 29.4 minutes while maintaining sensitivity of 0.565. Default options maintain a balance between execution speed and search sensitivity but can be tuned for specific applications as required.

### Domain recognition in multi-domain proteins

To investigate multi-domain search sensitivity, a benchmark dataset of AFDB^6^ multi-domain structures was assigned SCOPe fold classifications using SPfast. We evaluated the ability of each search method to identify multidomain partner proteins with a single overlapping domain (Figure 1B). The local alignment methods could retrieve partially overlapping pairs with 71.9%, 77.0% and 69.6% recall at the top-1 rank for SPalign, SPfast and foldseek (-e inf --max-seqs 200) respectively. Performance rose to 91.8%, 93.2% and 80.8% when considering matches within the top-25 structures. In this setting, there was a small benefit to SPfast re-ranking of the top 200 foldseek hits (foldseekSP) leading to a top-1 recall of 71.0% and top-25 recall of 81.7% respectively. The global alignment method, TMalign, was almost completely unable to recover partial structure matches, and identified only 42.9% of matched structures at the top-1 rank.

Surprisingly, SPscore optimization with SPalign underperformed SPfast in this benchmark due to systematic differences between experimental structures and predicted models. AFDB models contain isolated non-globular regions (often flexible, disordered loops predicted with low confidence) which can lead to disproportionately high scores between un-related structures when using a local alignment normalization strategy. Foldseek combats this problem by masking low complexity regions. Similarly, in SPfast, isolated residues are trimmed to avoid low-complexity regions based on the number of non-local neighbour residues. While a few low-complexity matches pollute the top ranks using SPalign, performance parity with SPfast is mostly restored when considering recall of partner proteins in the top-25 structures.

### Fold recognition of designed proteins

Current methods for *de novo* protein design generate backbone coordinates before assigning compatible amino acid sequences in a two-stage design process^20^. Here we evaluated the sensitivity of various structure-search methods using synthetic query structures designed by RFdiffusion to mimic SCOPe domain topologies without a corresponding amino acid sequence. The original foldseek (-s 9.5 --max-seqs 2000 -e 10 --comp-bias-corr 0 --mask 0 --alignment-type 0) scoring function was not optimized for backbone structure search and performed poorly in the absence of amino acid identities (median sensitivity of 0.273). In 2023, the foldseek scoring function was re-optimized for backbone-only structures leading to a dramatic performance improvement. However, the median sensitivity of foldseek-2023 (0.500) still falls far short of SPalign (0.643) and SPfast (0.667). Similarly, when we re-rank the top foldseek hits with SPfast (foldseekSP) the median sensitivity is significantly improved to 0.625.

We investigated the impact of increasing the degree of distortion introduced by the noising/denoising steps during the design process (Supplementary Figure S2). Performance for all methods was degraded by increasing the number of noise steps – likely reflecting the fact that the original SCOPe labels were not always appropriate for the sampled synthetic structures in the high-noise regime. Surprisingly, foldseek performance slightly improved in the low-noise regime compared with natural structures and may benefit from idealised synthetic motifs which are more closely aligned with the 3Di state alphabet. Threading the parent sequence on to the designed structures also improved foldseek performance at the family and superfamily level but was found to decrease sensitivity at the fold level. Similarly, sequence information appeared to buffer the degradation of performance with increasing noise which may be problematic if designed proteins have adversarial inconsistency between sequence and structure representations.

### HOMSTRAD alignment accuracy

We also evaluated the ability of each method to reproduce manually curated alignments from the HOMSTRAD^21^ database (Figure 2A). SPscore optimization was found to provide more accurate alignments than those produced by foldseek. However, TMalign was found to produce the most accurate alignments of all previously available methods. By default, SPscore was designed to operate with no penalty for alignment gaps which led to the inclusion of several isolated pairs which are unlikely to be representative of a true evolutionary relationship (Figure 2D). Introducing a gap open penalty of up to 0.5 (consistent with TMalign) dramatically improved alignment accuracy by eliminating these isolated pairs. However, improved alignment accuracy was accompanied by a small drop in search sensitivity as high-quality local matches (SCOPe false positives) out-ranked lower fidelity, domain-level hits (SCOPe true positives). SPscore-based methods relied on the extraneous matchings to dramatically increase the effective normalization length and partially suppress the score of high-fidelity alignments with low coverage.

**Figure 2.**
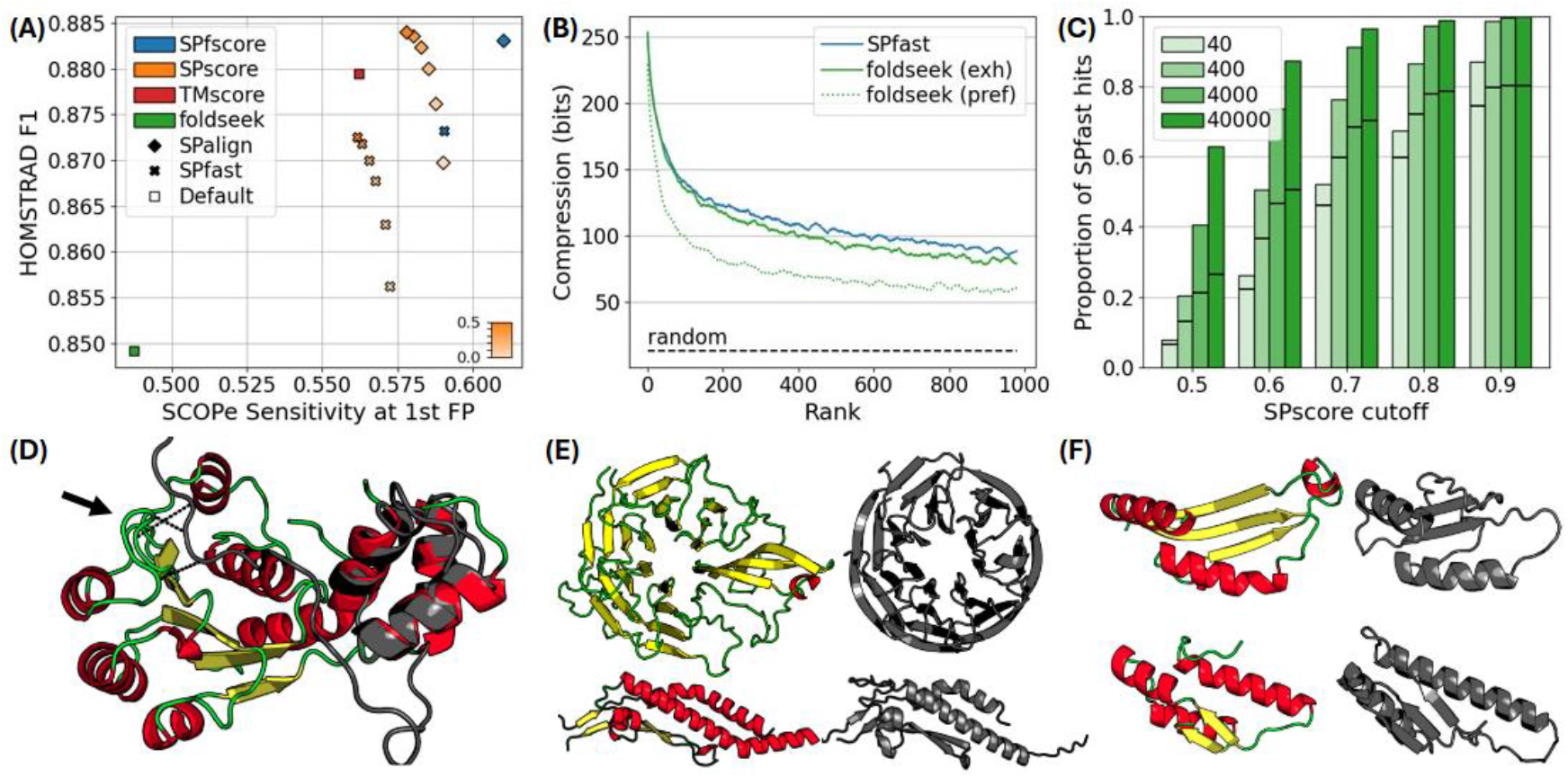
(A) Performance comparison of search sensitivity (SCOPe superfamily-level) and alignment accuracy while varying SPscore gap open penalty. (B) Pairwise alignment compression of top-ranking hits from AFDB clusters averaged over 100 random queries. The random baseline represents the average compression using an all-against-all comparison of the 100 queries. (C) Proportion of AFDB hits (defined by an SPscore cutoff) identified in the top-ranking structures by foldseek (--exhaustive-search). The solid horizontal lines indicate the results when using the foldseek prefilter at the indicated number of maximum sequences (40-40,000). (D) SPfast alignment between d1a5ta2 (color) and d1vg5a_ (gray) as representatives of different SCOPe folds with spurious aligned residues indicated. (E) Structure hits identified by SPfast but not found within the top 40,000 structures identified with the foldseek prefilter (upper) A0A4V0HUP4 – G8RV33 (rank: 203) and (lower) A0A7C1N324 – A0A7C3KYU8 (rank: 5). (F) Structure pairs identified by SPfast but not identified in the top 40,000 hits after exhaustive foldseek search. (upper) A0A832BBQ6 – A0A485P0T7 (rank: 102) and (lower) A0A3S3QK05 – A0A4P6JIK1 (rank: 507). Structures are trimmed to the aligned regions for visual clarity.

To overcome the trade-off between alignment accuracy and search sensitivity, we re-optimized SPscore parameters (d_0_, gap_open penalty and α) to control the balance between alignment coverage and fidelity. We found that this score re-parameterization significantly improved performance of both alignment accuracy and search sensitivity such that SPfast was equally sensitive with the original SPalign while also improving alignment accuracy and maintaining a >100x increased throughput. Combining the SPfast-optimized parameters with the original SPalign optimization algorithm also improved the performance of SPalign to achieve a new state-of-the-art performance for both HOMSTRAD alignment accuracy and SCOPe search sensitivity (Figure 2A).

### AFDB-clusters search sensitivity

To evaluate performance on a large predicted structure database we randomly selected 100 model structures from the AFDB-clusters^10^ dataset. These structures were searched against the full set of 2.3M AFDB-cluster representatives. In the absence of curated structural classifications, we used an objective, reference-free evaluation based on the degree of information compression afforded by pairwise structure alignments of top-ranked hits^22^. The very top-ranked foldseek candidates were found to have comparable quality to the top-ranked candidates proposed by SPfast (consistent with the family-level evaluation in the SCOPe benchmark). However, SPfast showed a sustained advantage in average compression for each position across the first 1000 ranks. Running foldseek with a permissive prefilter (fast: --max-seqs 40000 -e inf) was greatly improved by disabling the prefilter entirely (opt: --exhaustive-search) but was still unable to match the performance of SPfast (Figure 2B). SPfast search using the original SPscore parameters also provided a small improvement over foldseek in exhaustive mode (Supplementary Figure S3). Notably, the quality of retrieved structures by all methods was significantly above random pairs, indicating the presence of shared structural motifs even beyond the 1000^th^ rank.

We also investigated the potential to utilize foldseek to prefilter AFDB-clusters database prior to SPfast re-ranking. On average, 80% of extreme high-quality SPfast hits (SPscore > 0.9) could be identified within the first 400 structures ranked by foldseek bit score (Figure 2C). However, the remaining high-quality hits were not able to be identified even within the first 40,000 structures due to the nature of the *k*-mer based prefilter. We have highlighted several examples of top-ranking SPfast structure matches which were not identified by foldseek in the first 40,000 ranks (Figure 2E). For example, A0A4V0HUP4 is characteristic of a β-propeller domain with clear visual similarity to G8RV33 (rank 203 by SPfast). However, when using A0A4VHUP4 as a query, G8RV33 does not pass the foldseek prefilter and is not identified as a structural match. When disabling the prefilter with exhaustive search, similarity is identified and G8RV33 is recovered at rank 512. We have also highlighted a few examples of SPfast hits that were not identified by foldseek even in exhaustive mode (Figure 2F). These pairs display clear geometric similarity despite being poorly ranked by foldseek.

Finally, we highlight an example query (A0A6V8F1I9) to demonstrate the advantage of the global geometric approach employed by SPfast compared with the local structure alphabet employed by foldseek (Figure 3). A0A6V8F1I9 contains a repeated ββα structural motif where the helices rest on top of a conserved β-sheet. Based on the local structural neighbourhood, A0A017TCA9 contains the same segment sequence as A0A6V8F1I9 but arranged with a distinct global topology. Due to the compatible structural alphabet states (Figure 3E), foldseek ranks A0A017TCA9 highly (rank 386) at the expense of true geometric matches (eg A0A3D1EE05 – rank 2,232) which are correctly identified by SPfast (rank 10). In this example, global geometric information is critical to discriminate true structure matches from structure-state decoys.

**Figure 3.**
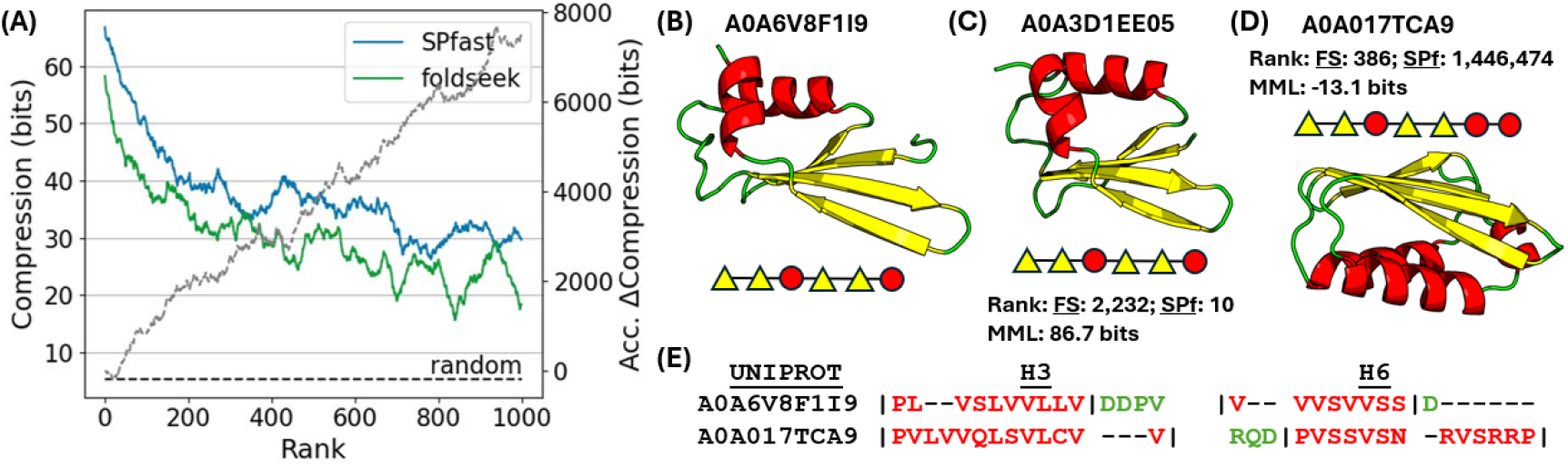
Characteristic example (A0A6V8F1I9) demonstrating the benefit of geometric search compared with a local structure alphabet. (A) Pairwise alignment compression of top-ranking hits from AFDB-clusters for A0A6V8F1I9 query (left) and accumulated compression advantage of SPfast-opt over foldseek-exhaustive (right). (B) A0A6V8F1I9 structure and segment diagram. (C) Structure and segment diagram of A0A3D1EE05 (D) Structure and segment diagram of A0A017TCA9. (E) foldseek 3Di encoding and alignment of helical segments from A0A6V8F1I9 and A0A017TCA9.

### AFDB dark clusters annotation

To demonstrate the potential utility of SPfast for highly sensitive annotation of protein functions, we extracted 46,826 high-quality (pLDDT>90) ‘dark’ clusters from the AFDB-clusters database. We used SPfast to search these uncharacterized proteins against the set of 2.3M AFDB-clusters representatives (more than 100B pairwise comparisons). Using SPfast, 25.2% of the dark clusters could be mapped to a high-complexity PFAM clan^23^. Using a foldseek cutoff of log_10_0.5, 31.9% of the annotated proteins shared at least 1 PFAM clan annotation in agreement with SPfast. An additional 45.2% of the annotated structures were uniquely annotated by SPfast while only 12.6% were uniquely annotated by foldseek – highlighting the complementarity of the 2 methods. The remaining 10.4% of proteins were annotated by both methods but with conflicting disjoint clan labels (Figure 4A). In addition, for each annotated protein, SPfast can recognize structural similarity to a larger number of PFAM clans.

**Figure 4.**
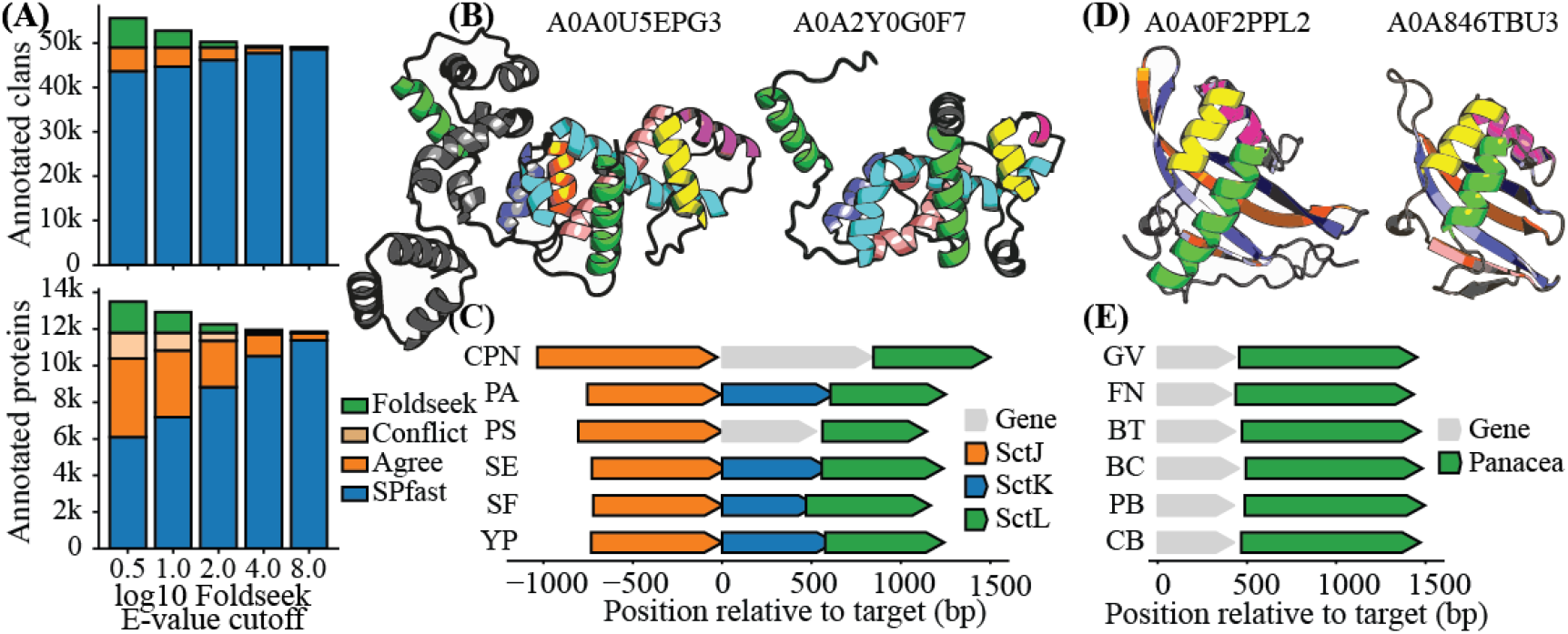
(A) Number of uncharacterized proteins from AFDB-clusters (pLDDT>90) that could be mapped to proteins with PFAM annotations based on structural similarity and the number of corresponding PFAM clan annotations. (B) Structure of uncharacterized protein A0A0U5EPG3 compared with A0A2Y0G0F7 which is annotated as a member of the Type III secretion system (SctK). (C) SctJKL genomic island in *Candidatus Protochlamydia naegleriophila* (CPN: GCF_001499655.1), *Pseudomonas aeruginosa PA01* (PA: GCA_000006765.1), *Pseudomonas syringae* (PS: GCA_041154885.1), *Salmonella enterica* (SE: GCA_019026085.1), *Shigella flexneri* (SF: GCA_003719775.1), *Yersinia pestis* (YP: GCA_003798345.1) assemblies. (D) Structure of uncharacterized protein A0A0F2PPL2 compared with A0A846TBU3 which is annotated as a Type II toxin. (E) Representative genomic context of structures from the A0A0F2PPL2 cluster including those from *Gardnerella vaginalis* (GV: GCA_001546485.1), *Fusobacterium necrophorum BFTR-2* (FN: GCA_000691725.1), *Bacillus thuringiensis serovar mexicanensis* (BT: GCA_002146325.1), *Bacillus cereus* (BC: GCA_002560615.1), *Peptococcaceae bacterium BRH_c4b* (PB: GCA_000961595.1), *Clostridiaceae bacterium* (CB: GCA_003485415.1) assemblies.

**Figure 5.**
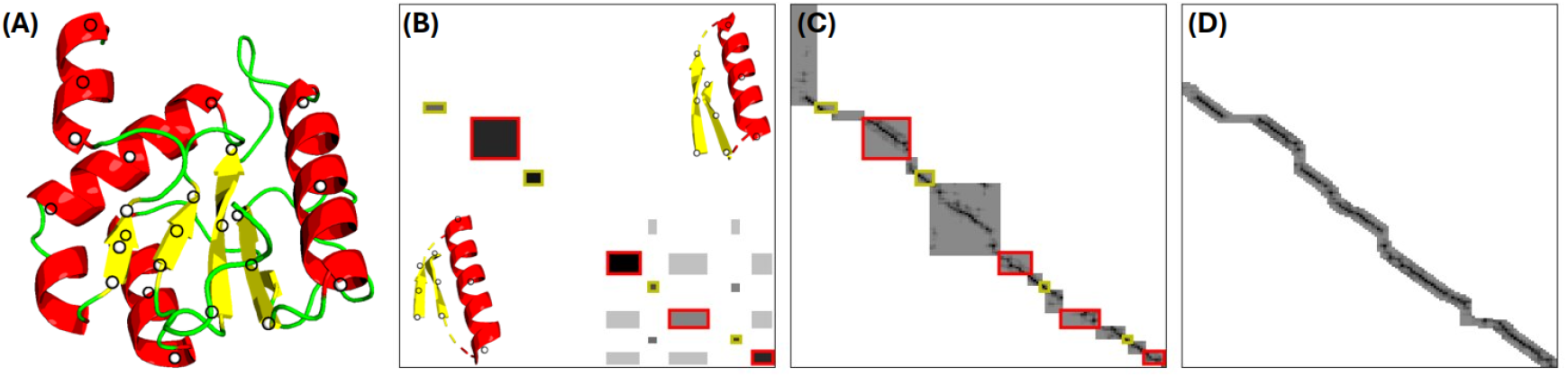
Pairwise alignment between d1w96a2 and d1a9xa3 SCOPe domains (A) d1a9xa3 domain with idealized segment-level keypoints. (B) A 3seg seed pair is used to superimpose the structures and produce an SPscore pairwise distance matrix between representative pseudo-atom coordinates. This score matrix is used to generate a segment-level sequence alignment. (C) A block-sparse all-atom distance matrix is constructed and constrained by the initial segment-level alignment. (D) A final all-residue refinement is conducted by exploring a local window of size 5 around the preliminary alignment.

As an example, the uncharacterized cluster, A0A0U5EPG3, is represented primarily by structures extracted from genomes in the *chlamydiota* phylum. This cluster demonstrated structural similarity to members of several PFAM clans (FliG, YscK, OrgA_MxiK, HrpB4, T3SS_LEE_assoc) which all have functions related to a type III secretion system (T3SS) adaptor protein responsible for connecting the sorting complex to the M ring^24^. These families correspond to the SctK gene under the unified T3SS nomenclature^25^ (Figure 4B). However, the corresponding gene in *chlamydiota* has not been previously reported despite several recent reviews^26,27^. T3SS is prevalent in pathogenic gram-negative bacteria and components are often found clustered in conserved genomic islands. A0A0U5EPG3 in *Candidatus Protochlamydia naegleriophila* is flanked by the SctJ and SctL genes which is consistent with the genomic context of the SctK gene in phyla where it has been annotated (Figure 4C). The combination of molecular and syntenic similarity provides orthogonal support for a shared functional role of the uncharacterized proteins in the A0A0U5EPG3 cluster with the well-characterized T3SS SctK gene. Furthermore, AlphaFold3 (AF3) predicts a high-confidence ternary structure between A0A0U5EPG3, the second forkhead association (FHA2) domain of the inner membrane ring protein (SctD) and the N-terminal domain of the cytosolic sorting platform (SctQ), consistent with the reported role of the SctK gene in other organisms^28^. Interestingly, in the AF3 model, the SctQ-A0A0U5EPG3 interface is mediated by the C-terminal lobe which is unique to the *chlamydiota* phylum.

Similarly, A0A0F2PPL2 from *Peptococcaceae bacterium BRH_c4b* is representative of a cluster of uncharacterized proteins dominated by members from *Bacillota* genomes. SPfast identifies A0A846TBU3 as the top structural match which is annotated as a Type II toxin from the MqsR^29^ family (Figure 4D). Inspecting the genomic context of the A0A0F2PPL2 cluster reveals that almost every member can be found immediately adjacent to a MqsA-Panacea^30^ antitoxin protein (Figure 4E). Based on the AF3 model complex, the Zn^2+^-binding domain forms a high-confidence interface with the putative toxin, providing additional support for a toxin-antitoxin functional assignment. Furthermore, A0A0F2PPL2 contains a conserved Tyrosine (Y109) in the putative RNA-binding groove which has been identified as a critical catalytic residue in the *B. fungorum* MsqR endoribonuclease toxin^30^ adding further evidence to suggest a shared functional role.

## Discussion

Since the release of AlphaFold2^31^, the number of protein structures available in public databases^6,8^ has been increased by three orders of magnitude compared with what was previously available in the PDB^32^. Similarly, the emergence of new predictive models for *de novo* design^20^, increased availability of dynamic trajectories^33^ and alternative models for structure prediction^34,35^ will continue to drive a surge in protein structure data. This sustained data influx motivates the development of new bioinformatic tools that can scale to meet the rising demand. In this work we have introduced a new method for geometric structure comparison (SPfast) which achieves state of the art search sensitivity and improves alignment accuracy over a range of performance benchmarks. While we do not achieve the same speed as the tokenized alignment implemented in foldseek, we argue that the speed of SPfast is sufficient for practical use – particularly for applications that prioritize high sensitivity search such as supporting biological experiments which are conducted over long timeframes.

Compared with traditional methods, SPfast achieves remarkable acceleration by 1) superimposing only a minimal set of structure fragments to define alignment seeds 2) filtering candidate structures with an efficient segment-level alignment and 3) using the segment-level alignment as a constraint to produce a block-sparse distance matrix. On average, segment-guided superpositions are better quality than those produced by arbitrary contiguous fragments and better facilitate the discrimination of structural similarity at early stages of the alignment. Similarly, unlike tokenized substitution matrices, which create dense score matrices, geometric matches are necessarily unique since atomic coordinates from the same protein cannot be overlapping. As a result, the segment-level constraints greatly reduce the time required to compute all-residue distances matrices which are re-evaluated many times during ICP optimization.

In extremely rare cases, the minimal set of segment-guided seeds do not produce a high-quality superposition which degrades the quality of the final alignment. As future work, we are exploring options to refine the set of seed comparisons (eg by utilizing a segment-level 3di state rather than simplistic secondary structures) which may further improve both sensitivity and computational efficiency. Similarly, we are investigating options to rescue poor initial superpositions by better sampling candidate alignments (eg by using pre-computed neighbour graphs to explore alternative matchings). Finally, the current SPfast implementation does not take advantage of SIMD intrinsics to vectorize alignment and superposition (as implemented in foldseek). While SIMD operations can be of limited value for sparse data, the block-sparse nature of the SPfast algorithm would facilitate efficiency gains enabled by SIMD operations. These proposed refinements represent several viable strategies for further acceleration and improved sensitivity.

As a proof of principle, we have used SPfast to identify candidate annotations for a large database of ‘dark’ proteins which lack existing functional information. From this dataset, we have highlighted two examples of SPfast-based annotations which are supported by independent *in silico* evidence. The identification of a putative SctK gene in *Chlamydiota* provides new insights into a major virulence factor responsible for modulating host cell biology in a highly relevant human pathogen. Similarly, the annotation of new bacterial toxins sheds light on key cellular regulators. These putative annotations are strongly supported by synteny relationships and high confidence AlphaFold3 interactions. However, these examples are by no means comprehensive. For example, the dataset is also enriched with annotations of endonucleases with HTH domains and key conserved catalytic residues, as well as putative metalloproteases with HEXXH motifs supporting the SPfast-based assignments. Finally, numerous structural relationships have also been identified by SPfast without clear *in silico* support for the resulting functional assignments.

## Methods

### Secondary structure assignment

Secondary structure states are assigned based on the union of the permissive Cα-distance-based definition currently utilized by TMalign^14^/SPalign^15^ as well as the DSSP definition based on electrostatic interactions^36^. Protein structure models utilize the DSSP definition only (without Cα-based definition) to avoid an artifact which causes long disordered loops to be classified as strands. Segment breaks are introduced at residues classified as geometric turns by DSSP to maintain equivalence between secondary structure segments of the same type (ie long twisted beta strands are not geometrically comparable to short straight strands). We enforce a minimum segment length of 6 for helices and 3 for strands, as shorter segments appear to be less evolutionarily conserved and can obscure seed matches between contiguous segments.

### Secondary structure prefilter

To reduce the total number of required comparisons during SCOPe database search, we prefilter the library to avoid aligning proteins from clearly un-related protein classes. We utilize a sequential representation of secondary structure segments by assigning helix and sheet labels to corresponding fragments. Structures are aligned based on the resulting segment-state sequence and further processed if the alignment score reaches a critical score threshold. Structures that do not pass the preliminary prefilter are removed from the comparison set. This prefilter was used for single domain datasets (such as SCOPe) whereby the alignment score can be normalized by the total number of segments in the structures.

### Representative key-point generation

We reduce the all-residue representation of protein structure to a sequence of secondary structure segments. To compensate for the varied segments lengths, we extract 3 representative key-points including the centroid and two terminal residues. Pseudo-atom coordinates are idealized by projecting true atomic coordinates on to the first principal component of each segment to minimize the contribution of residue periodicity.

### Alignment score

In this work, we use the SPscore alignment objective reported previously^15^.

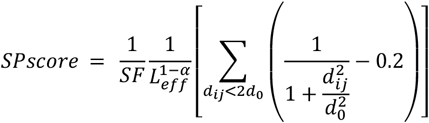

Where d_ij_ is the distance between Cα atoms of aligned residues and d_0_ and α are free parameters optimized in prior work. SPscore is normalized by an effective length (L_eff_) which is dependent on the residue-level alignment. Core aligned residues are defined by correspondences with pairwise distances <2d_0_. L_eff_ is defined by the number of core residues combined with the average number of neighbouring residues within a 3d_0_ window. SF is a purely cosmetic scaling factor which ensures that the typical fold discrimination cutoff falls approximately in a familiar range around 0.5. The coarse-grained, segment-level alignment optimizes an objective with the same form as SPscore but averaged over the three representative pseudo-atoms.

### Optimization algorithm

We initially superimpose sets of 3 contiguous segments (3Segs) extracted from parent structures and prune seed superpositions with local fragment RMSD greater than a critical threshold. Successful seeds are extended to evaluate domain-level similarity using a coarse-grained, segment-level alignment optimizing the SPscore objective (Figure 3B). Subsequently, the process is repeated with all possible combinations of 2Segs where the 2Seg definition is extended to include skip-pairs (ie pairs that are separated by an interloping segment). To minimize redundant evaluations, 3Segs that pass the initial quality filter are decomposed into the set of possible 2segs and excluded from re-analysis. Based on the segment-level similarity score, the optimum seed is selected to generate residue-level correspondences. A block-sparse Smith-Waterman alignment is employed to identify a residue-level alignment constrained by initial segment matchings (Figure 3C). A final iterative refinement stage is conducted which involves evaluating potential alignment pairs in a 5-residue window around the preliminary alignment (Figure 3D). Superpositions are updated based on candidate alignments at each iteration. After convergence, final alignments are conducted with a default gap open penalty of 0.2.

### SCOPe benchmark

We evaluate SPfast in the SCOPe benchmark reported previously for the evaluation of foldseek^18^. Briefly, we generate all-against-all alignments between 11,211 non-redundant SCOPe domains with curated fold annotations. For each query, the remaining structures are ranked, and methods are evaluated based on their ability to recover proteins with the same structural classification at all 3 levels of the hierarchy (fold, superfamily, family). Performance is evaluated based on the sensitivity at the first-ranked false positive.

### Multi-domain benchmark

We collected representative structures from the AFDB-clusters dataset^10^ and conducted structure search against the SCOPe domains to assign SCOPe fold classifications based on SPfast structural similarity (SP>0.55, L_eff_>100, L_afdb_ > 1.2 x L_SCOPe_). We retained AFDB structures that could be assigned multiple structurally distinct SCOPe domains where structural similarity between domain archetypes was determined by SPalign (SP<0.45 between each assigned domain). Finally, we randomly selected two multi-domain AFDB structures for each qualifying SCOPe domain. We searched each structure against the complete dataset and evaluated performance based on the rank of the assigned partner for each domain. Structures that incidentally contained the same SCOPe classification (SP>0.45 to shared domain) but were not in the assigned pair were excluded during the evaluation.

### Designed protein benchmark

To generate diverse synthetic structures, we noised and de-noised SCOPe domains using the partial diffusion protocol from RFdiffusion^20,37^. We applied noise for 10 steps and then denoised the resulting distorted structure to produce a distinct, backbone-only approximation for each of the representative SCOPe domains. It was assumed that synthetic domains would have the same fold classification as the parent structure. Synthetic domains were used to search the SCOPe database of natural proteins and performance was evaluated at the superfamily level based on the sensitivity at the first false positive.

### HOMSTRAD alignment benchmark

The HOMSTRAD alignment benchmark^21^ includes manually curated alignments for 1032 protein families. We reproduced the alignment pairs used to evaluate foldseek by extracting the first and last family member from the 2022_Aug_1 release. Note that there is a slight difference between structures used in this work and those reported previously^18^ since the HOMSTRAD dataset is no longer publicly available and was reproduced from best efforts. Alignment performance was evaluated based on F1 score (harmonic mean of precision and recall) by comparing the set of alignment correspondences with the correspondences from the gold-standard reference alignments (https://wwwuser.gwdg.de/~compbiol/foldseek/). Geometric alignments included residue pairs within 5Å.

### AFDB-clusters database

To evaluate search performance on a large model dataset, we collected ∼2.3M representative structures from the AFDB-clusters^10^ database (https://afdb-cluster.steineggerlab.workers.dev/). Structure coordinates were extracted from the compressed foldcomp^38^ repository and then pre-processed to extract idealized secondary structure segments identified by DSSP. AFDB model structures contain regions of long extended loops predicted with low confidence which are likely to be intrinsically disordered. To gracefully handle these cases, we trimmed the structures to remove non-compact regions based on the number of non-local (i-j>8) contacts (distance<12Å). This procedure is similar to the low complexity masking employed by foldseek but has the added benefit of reducing the effective protein length to facilitate further accelerated search.

### Alignment compression

We used MMLigner^22^ to quantify the maximum degree of compression afforded by pairwise structural alignments as a reference-free evaluation of search sensitivity in the AFDB-clusters benchmark. Briefly, a Kent mixture model was used to encode internal pseudo-angles as a null model of independent protein structures. Theoretical compression was defined as the information saved by encoding structures jointly conditioned on the pairwise alignment compared with the null model. Compression at each rank was smoothed by averaging in a window of 10 to aid visual clarity in the figure (window of 100 for A0A6V8F1I9 case study).

### Foldseek

For consistency, unless otherwise stated we use the foldseek commit (aeb5e) and commands as reported in the original manuscript. Only time taken for the ‘prefilter’, ‘structurealign’ and ‘tmalign’ operations were considered when evaluating foldseek-based execution time. Commit ef4e9 was used to evaluate backbone only search (--comp-bias-corr 0 --mask 0 --alignment-type 0).

### Foldseek prefilter

FoldseekTM was developed to improve the sensitivity of foldseek by re-ranking top scoring candidates with an optimized version of TMalign. In this work, we extend this idea to produce foldseekSP which pre-filters candidate pairs using foldseek before re-ranking structures by SPfast alignment.

### TMalign

TMalign was downloaded from https://zhanggroup.org/TM-align/TMalign.cpp using the version updated on 2022/04/12. Consistent with prior work^18^, alignments were ranked by average TMscore using both the shorter and longer protein lengths. We found that this strategy produced the best results in the SCOPe benchmark.

### PFAM Annotations

PFAM annotations were downloaded from https://ftp.ebi.ac.uk/pub/databases/Pfam/releases/Pfam37.0/ and https://ftp.ebi.ac.uk/pub/databases/interpro/releases/101.0/. Annotations were filtered to ensure that the annotated length covered >60% of the reference family. After mapping to trimmed AFDB-clusters structures, annotations were filtered to remove low complexity structures with <5 helices or <4 mixed segments.

### AFDB dark clusters

We extracted high-quality (pLDDT>90) ‘dark’ clusters from the AFDB-clusters dataset which could not be mapped to PFAM domains. ‘Dark’ clusters were searched against the entire AFDB-clusters dataset with SPfast. Foldseek cluster similarity results from 2023-06-05 were downloaded fromhttps://afdb-cluster.steineggerlab.workers.dev/. PFAM annotations were assigned to dark clusters using high-scoring alignments with annotated cluster representatives. Structure matches were filtered such that queries contained at least 6 aligned helices or 5 aligned segments for mixed domains. Annotations were transferred to the query structure if aligned residues covered at least 70% of the PFAM domain from the reference structure. To match the permissive foldseek alignment lengths, SPfast alignments were evaluated by including residue pairs within 8Å.

### Genomic context

Genomic context was extracted using a modified version of GCsnap^39^. Briefly, UniProt IDs were mapped to GenkBank CDS ID using the UniProt id-mapping api. The GenBank ID was mapped to a GenBank assembly and the target protein genomic context was extracted using the NCBI Entrez api.

### AlphaFold3 structure prediction

Complex structures were predicted using the AlphaFold3 web server (https://alphafoldserver.com/). Full length sequences were trimmed to the expected interacting domains for visual clarity and to isolate direct interactions in the predicted aligned error (PAE) plot.

### Data availability

Pre-computed benchmark data is made available at https://spfast.tomlitfin.workers.dev/. The SPfast source code (including a PyMOL plugin and demonstrative colab notebook) is available from https://github.com/tlitfin/SPfast.

## Acknowledgement

We gratefully acknowledge the support of the Griffith University eResearch Service & Specialised Platforms Team and the use of the High-Performance Computing Cluster “Gowonda” to complete this research. TL is supported by a Griffith University Postgraduate Fellowship. MvI is supported by the National Health and Medical Research Council, Australia (NHRMC, ID 2009677 & GNT1196520). YZ is supported by Natural Science Foundation of China (Grant #:92370202) and the computing facility at Shenzhen Bay Laboratory. In addition, TL acknowledges Lenovo who provided a Thinkstation workstation to support this research. We also thank Professor Yuedong Yang for making the SPalign source code freely available.

## Notes

https://github.com/tlitfin/SPfast

https://spfast.tomlitfin.workers.dev/

### Competing Interest Statement

The authors have declared no competing interest.

### Summary of Updates

Minor adjustment to parameters used for function annotation in Figure 4. Updated methods to provide additional details.

## References

1. Seemann, T. Prokka: rapid prokaryotic genome annotation. Bioinformatics 30, 2068–2069 (2014).

2. Altschul, S. F., Gish, W., Miller, W., Myers, E. W. & Lipman, D. J. Basic local alignment search tool. J. Mol. Biol. 215, 403–410 (1990).

3. Eddy, S. R. Accelerated Profile HMM Searches. PLOS Comput. Biol. 7, e1002195 (2011).

4. Steinegger, M. & Söding, J. MMseqs2 enables sensitive protein sequence searching for the analysis of massive data sets. Nat. Biotechnol. 35, 1026–1028 (2017).

5. Weisman, C. M., Murray, A. W. & Eddy, S. R. Many, but not all, lineage-specific genes can be explained by homology detection failure. PLOS Biol. 18, e3000862 (2020).

6. Varadi, M. et al. AlphaFold Protein Structure Database: massively expanding the structural coverage of protein-sequence space with high-accuracy models. Nucleic Acids Res. 50, D439–D444 (2022).

7. Consortium, U. UniProt: the universal protein knowledgebase in 2023. Nucleic Acids Res. 51, D523–D531 (2023).

8. Lin, Z. et al. Evolutionary-scale prediction of atomic-level protein structure with a language model. Science (80-. ). 379, 1123–1130 (2023).

9. Pavlopoulos, G. A. et al. Unraveling the functional dark matter through global metagenomics. Nature 622, 594–602 (2023).

10. Barrio-Hernandez, I. et al. Clustering predicted structures at the scale of the known protein universe. Nature 622, 637–645 (2023).

11. Borujeni, P. M. & Salavati, R. Functional domain annotation by structural similarity. NAR Genomics Bioinforma. 6, lqae005 (2024).

12. Rodríguez del Río, Á. et al. Functional and evolutionary significance of unknown genes from uncultivated taxa. Nature 626, 377–384 (2024).

13. Cohen, G. E. ALIGN: a program to superimpose protein coordinates, accounting for insertions and deletions. J. Appl. Crystallogr. 30, 1160–1161 (1997).

14. Zhang, Y. & Skolnick, J. TM-align: a protein structure alignment algorithm based on the TM-score. Nucleic Acids Res. 33, 2302–2309 (2005).

15. Yang, Y., Zhan, J., Zhao, H. & Zhou, Y. A new size-independent score for pairwise protein structure alignment and its application to structure classification and nucleic-acid binding prediction. Proteins Struct. Funct. Bioinforma. 80, 2080–2088 (2012).

16. Zhang, C., Shine, M., Pyle, A. M. & Zhang, Y. US-align: universal structure alignments of proteins, nucleic acids, and macromolecular complexes. Nat. Methods 19, 1109–1115 (2022).

17. Needleman, S. B. & Wunsch, C. D. A general method applicable to the search for similarities in the amino acid sequence of two proteins. J. Mol. Biol. 48, 443–453 (1970).

18. van Kempen, M. et al. Fast and accurate protein structure search with Foldseek. Nat. Biotechnol. 42, 243–246 (2024).

19. Chandonia, J.-M., Fox, N. K. & Brenner, S. E. SCOPe: classification of large macromolecular structures in the structural classification of proteins-extended database. Nucleic Acids Res. 47, D475–D481 (2019).

20. Watson, J. L. et al. De novo design of protein structure and function with RFdiffusion. Nature 620, 1089–1100 (2023).

21. Mizuguchi, K., Deane, C. M., Blundell, T. L. & Overington, J. P. HOMSTRAD: a database of protein structure alignments for homologous families. Protein Sci. 7, 2469–2471 (1998).

22. Collier, J. H. et al. Statistical inference of protein structural alignments using information and compression. Bioinformatics 33, 1005–1013 (2017).

23. Mistry, J. et al. Pfam: The protein families database in 2021. Nucleic Acids Res. 49, D412–D419 (2021).

24. Soto, J. E., Galán, J. E. & Lara-Tejero, M. Assembly and architecture of the type III secretion sorting platform. Proc. Natl. Acad. Sci. 119, e2218010119 (2022).

25. Wagner, S. & Diepold, A. A Unified Nomenclature for Injectisome-Type Type III Secretion Systems BT - Bacterial Type III Protein Secretion Systems. in (eds. Wagner, S. & Galan, J. E.) 1–10 (Springer International Publishing, 2020). doi:10.1007/82_2020_210

26. Deng, W. et al. Assembly, structure, function and regulation of type III secretion systems. Nat. Rev. Microbiol. 15, 323–337 (2017).

27. Rucks, E. A. Type III Secretion in Chlamydia. Microbiol. Mol. Biol. Rev. 87, e0003423 (2023).

28. Muthuramalingam, M. et al. The Structures of SctK and SctD from Pseudomonas aeruginosa Reveal the Interface of the Type III Secretion System Basal Body and Sorting Platform. J. Mol. Biol. 432, 166693 (2020).

29. Sanchez-Torres, V., Kirigo, J. & Wood, T. K. Diverse physiological roles of the MqsR/MqsA toxin/antitoxin system. Sustain. Microbiol. 1, qvae006 (2024).

30. Ernits, K. et al. The structural basis of hyperpromiscuity in a core combinatorial network of type II toxin– antitoxin and related phage defense systems. Proc. Natl. Acad. Sci. 120, e2305393120 (2023).

31. Jumper, J. et al. Highly accurate protein structure prediction with AlphaFold. Nature 596, 583–589 (2021).

32. Berman, H. M. et al. The Protein Data Bank. Nucleic Acids Res. 28, 235–242 (2000).

33. Tiemann, J. K. S. et al. MDverse: Shedding Light on the Dark Matter of Molecular Dynamics Simulations. Elife 2023.05.02.538537 (2023). doi:10.7554/elife.90061.1

34. Stahl, K., Graziadei, A., Dau, T., Brock, O. & Rappsilber, J. Protein structure prediction with in-cell photo-crosslinking mass spectrometry and deep learning. Nat. Biotechnol. 41, 1810–1819 (2023).

35. Abramson, J. et al. Accurate structure prediction of biomolecular interactions with AlphaFold 3. Nature 630, 493–500 (2024).

36. Kabsch, W. & Sander, C. Dictionary of protein secondary structure: Pattern recognition of hydrogen-bonded and geometrical features. Biopolymers 22, 2577–2637 (1983).

37. Vázquez Torres, S. et al. De novo design of high-affinity binders of bioactive helical peptides. Nature 626, 435–442 (2024).

38. Kim, H., Mirdita, M. & Steinegger, M. Foldcomp: a library and format for compressing and indexing large protein structure sets. Bioinformatics 39, btad153 (2023).

39. Pereira, J. GCsnap: Interactive Snapshots for the Comparison of Protein-Coding Genomic Contexts. J. Mol. Biol. 433, 166943 (2021).

