## Supplementary Material for "Ultra-fast and highly sensitive protein structure alignment with segment-level representations and block-sparse optimization"

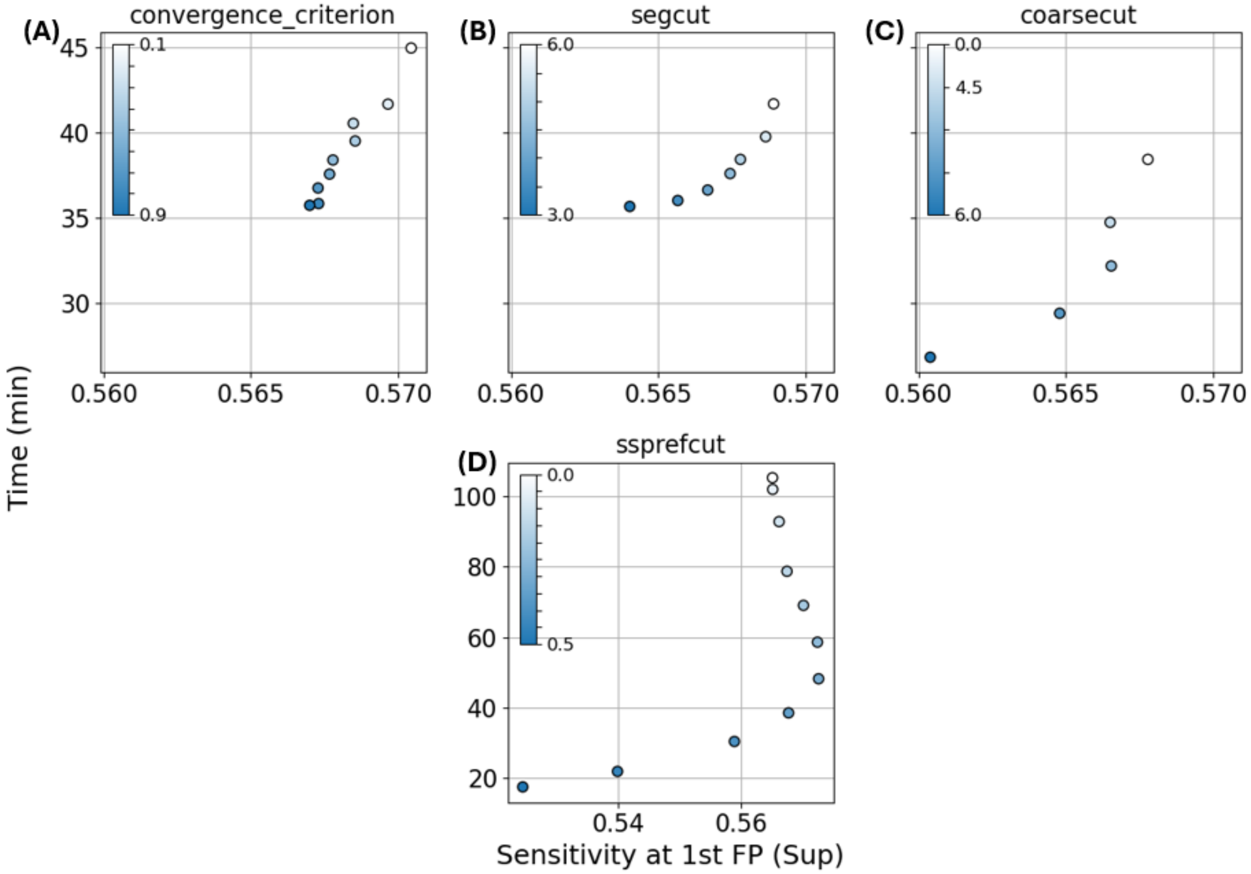


Figure S1 Mean superfamily sensitivity at the first false positive for SCOPe domains and corresponding execution time for 11,211 x 11,211 all-by-all comparisons on an Intel Xeon E5-2670 @ 2.60GHz 16-core CPU. (A) Performance variation when adjusting the convergence criterion (default: 0.5) for the iterated optimization. (B) Performance variation when adjusting the segment seed RMSD cutoff for seed pruning (default: 5.0). (C) Performance variation when adjusting the segment-level prefilter score cutoff (default: 0). (D) Performance variation when adjusting the secondary structure prefilter score cutoff (default: 0.35).

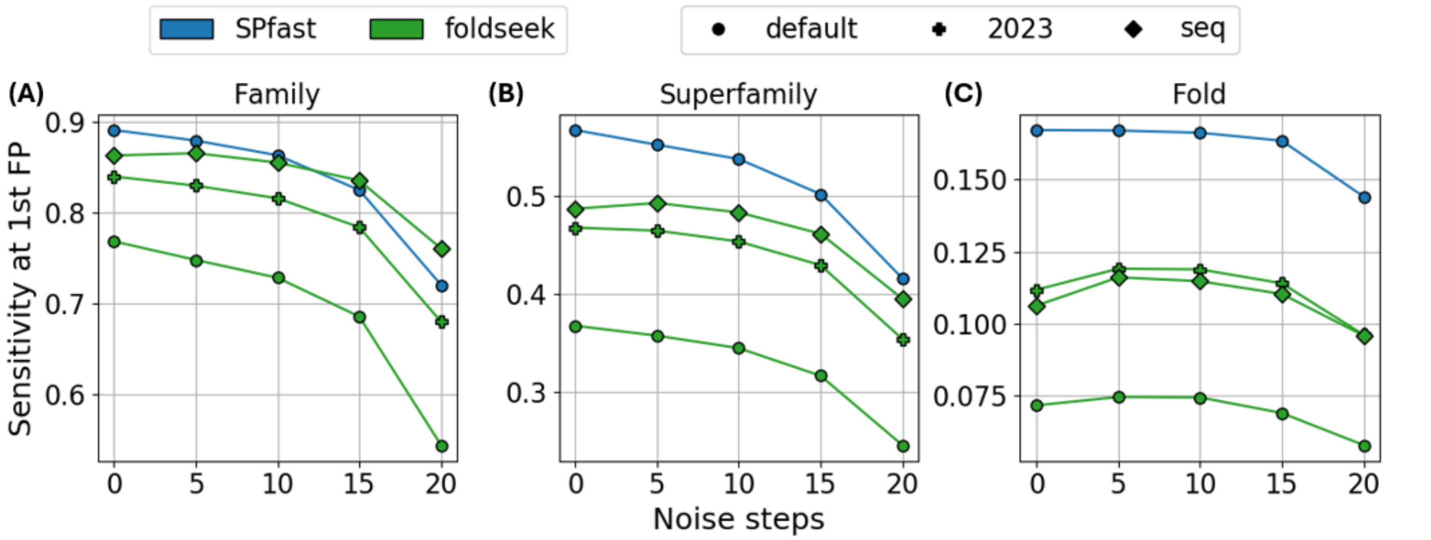


Figure S2 Mean sensitivity at the first false positive at (A) family, (B) superfamily and (C) fold level for synthetic structures generated using the RFdiffusion partial denoising protocol on native SCOPe domains. foldseek performance is evaluated using backbone-only structures with the default and 2023 implementation as well as by threading the parent SCOPe sequence on to the synthetic designed structure (seq).

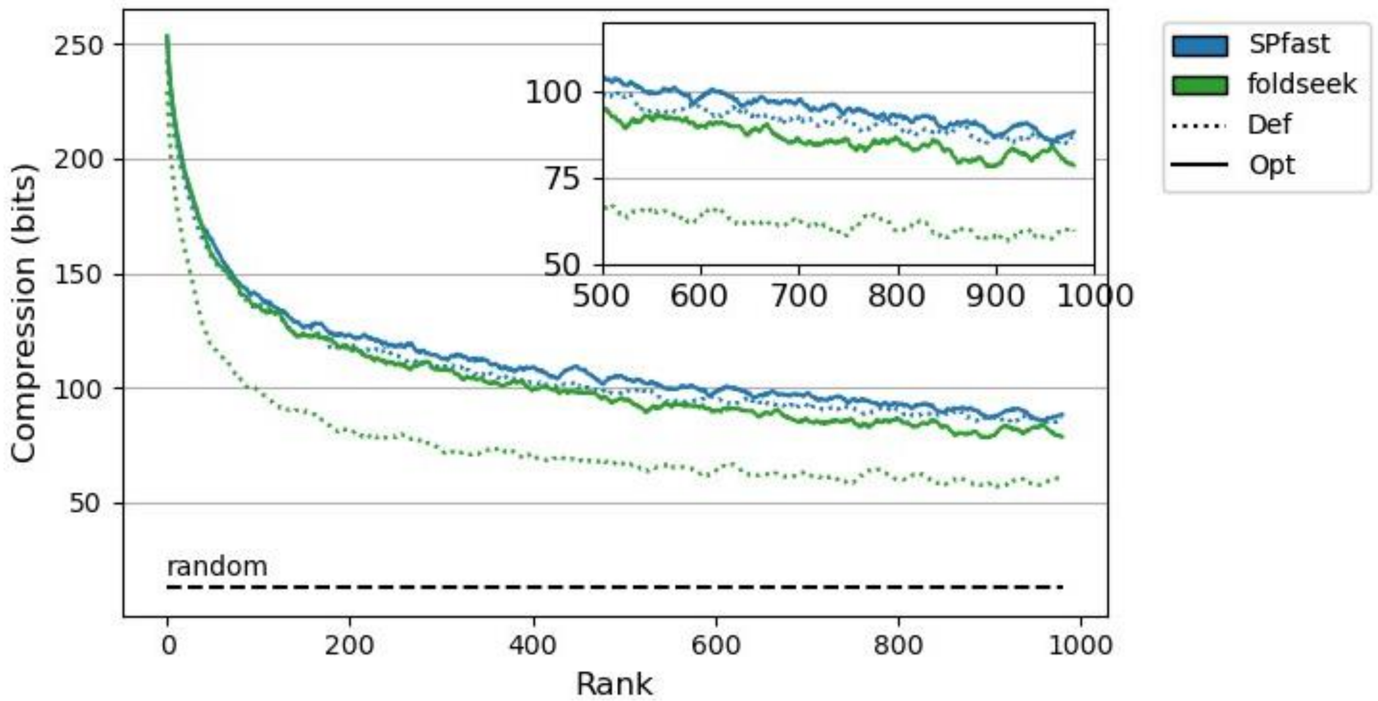


Figure S3 Pairwise alignment compression of top-ranking hits from AFDB-clusters averaged over 100 random queries. SPfast is run with default (Def) and optimized (Opt) parameters and foldseek is run with “--max-seqs 40,000 -e inf” (Def) and “--exhaustive-search -e inf” (Opt) arguments.

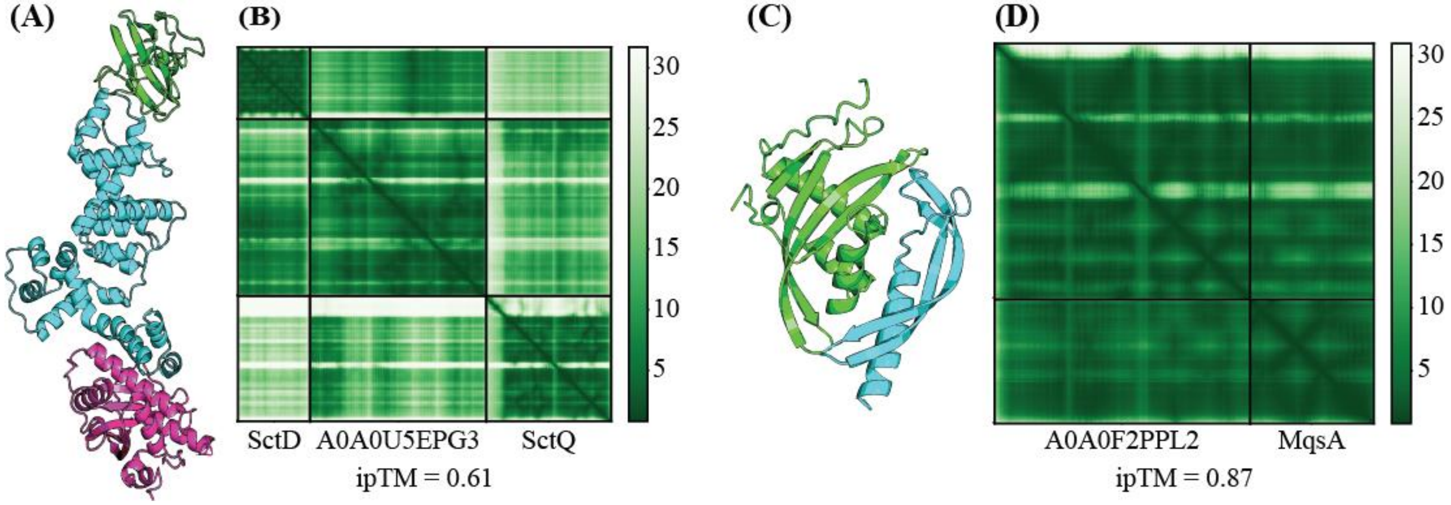


Figure S 4 (A) AlphaFold3 prediction for complex structure between the FHA2 domain from the SctD gene (172-285), A0A0U5EPG3 and the N-terminus of SctQ (1-199) all extracted from the GCF_001499655.1 *Candidatus Protochlamydia naegleriophila* assembly (B) PAE plot of the AlphaFold3 prediction for FHA2 domain from SctD, A0A0U5EPG3 and the N-terminus of SctQ. (C) AlphaFold3 prediction for complex structure between A0A0F2PPL2 and the $Zn^{2+}$-binding domain of the MqsA gene (1-69) immediately downstream of A0A0F2PPL2 in the GCF_000961595.1 *Peptococcaceae bacterium BRH_c4b* assembly. (D) PAE plot of the AlphaFold3 prediction for A0A0F2PPL2 and the $Zn^{2+}$-binding domain from MqsA.
